# Using the basic reproduction number to assess the risk of transmission of lumpy skin disease virus by biting insects

**DOI:** 10.1101/602292

**Authors:** Simon Gubbins

**Affiliations:** The Pirbright Institute, Ash Road, Pirbright, Surrey GU24 0NF, U.K.

**Keywords:** cattle, epidemiology, transmission model, Bayesian methods, uncertainty analysis, sensitivity analysis

## Abstract

In recent years, lumpy skin disease virus (LSDV) has emerged as a major threat to cattle outside Africa, where it is endemic. Although evidence suggests that LSDV is transmitted by the bites of blood sucking arthropods, few studies have assessed the risk of transmission posed by particular vector species. Here this risk is assessed by calculating the basic reproduction number (*R*_0_) for transmission of LSDV by five species of biting insect: the stable fly, *Stomoxys calcitrans*, the biting midge, *Culicoides nubeculosus*, and three mosquito species, *Aedes aegypti*, *Anopheles stephensi* and *Culex quinquefasciatus*. Parameters relating to mechanical transmission of LSDV were estimated using new analyses of previously-published data from transmission experiments, while vector life history parameters were derived from the published literature. Uncertainty and sensitivity analyses were used to compute *R*_0_ for each species and to identify those parameters which influence its magnitude. Results suggest that *S. calcitrans* is likely to be the most efficient at transmitting LSDV, with *Ae. aegypti* also an efficient vector. By contrast, *C. nubeculosus*, *An. stephensi*, and *Cx. quinquefasciatus* are likely to be inefficient vectors of LSDV. However, there is considerable uncertainty associated with the estimates of *R*_0_, reflecting uncertainty in most of the constituent parameters. Sensitivity analysis suggests that future experimental work should focus on estimating the probability of transmission from insect to bovine and on the virus inactivation rate in insects.

## 1 Introduction

Lumpy skin disease (LSD) is an important transboundary disease of cattle and is caused by lumpy skin disease virus (LSDV). Historically, LSD outbreaks have largely been confined to Africa (EFSA Panel on Animal Health and Welfare, 2015), where the disease contributes to rural poverty and food insecurity (Tuppurainen & Oura, 2012; Molla, de Jong, Gari & Frankena, 2017). In recent years, however, LSDV has emerged as a major threat to cattle outside of Africa. In 2012 it spread to the Middle East, through Israel and the Lebanon, reaching Turkey in 2013 (Alkhamis & VanderWaal, 2016). In 2015, the first cases of LSD were reported in Greece (Tasioudi et al., 2016) and the virus subsequently spread to much of the Balkans (Mercier et al., 2018).

Evidence from the field and from experiments indicates that transmission of LSDV by most direct and indirect routes is inefficient (Weiss, 1968; Carn & Kitching, 1995). The principal transmission route for LSDV is believed to be via the bites of blood sucking arthropods (Tuppurainen & Oura, 2012; EFSA Panel on Animal Health and Welfare, 2015). This has been shown experimentally for *Aedes aegypti* mosquitos (Chihota, Rennie, Kitching & Mellor, 2001) and for *Rhipicephalus appendiculatus* male ticks (Tuppurainen et al., 2013). The stable fly, *Stomoxys calcitrans*, has been incriminated as a potential vector in Israel by comparing its seasonal abundance with the seasonality of LSD cases (Kahana-Sutin, Klement, Lensky & Gottlieb, 2017) and because of its ability to transmit the closely related capripox virus between sheep and goats (Kitching & Mellor, 1986; Mellor, Kitching & Wilkinson, 1987). The potential for the biting midge, *Culicoides nubeculosus*, and two other mosquito species, *Culex quinquefasciatus* and *Anopheles stephensi*, to transmit LSDV has also been assessed, though for these three species attempts at transmission were unsuccessful (Chihota, Rennie, Kitching & Mellor, 2003).

In this study, the risk of transmission of LSDV posed by different insect species is explored by estimating the basic reproduction number for five species of biting insect (*S. calcitrans*, *C. nubeculosus*, *Ae. aegypti*, *An. stephensi* and *Cx. quinquefasciatus*). These species were selected as their potential role in LSDV transmission has been investigated previously (Chihota, Rennie, Kitching & Mellor, 2001, 2003) and they represent insect species (or at least genera) that are relevant to Europe. The basic reproduction number, denoted by *R*_0_, is the “average number of secondary cases arising from the introduction of a single infected individual into an otherwise susceptible population” (Diekmann & Heesterbeek, 2000). An outbreak can occur only if *R*_0_>1 and, consequently, *R*_0_ provides a means to assess the risk posed by each vector species.

Using a transmission model, an expression for *R*_0_ is derived that shows how it relates to the underlying transmission processes in insect vectors and cattle. These constituent parameters are then estimated using data from the published literature. In particular, data from transmission experiments involving the five putative vector species (Chihota, Rennie, Kitching & Mellor, 2001, 2003) are re-analysed using Bayesian methods to quantify the uncertainty in parameters relating to mechanical transmission. In addition, the latent and infectious periods for LSDV are estimated from the outcome of challenge experiments (Tuppurainen, Venter & Coetzer, 2005; Babiuk et al., 2008), again using Bayesian methods. Finally, uncertainty and sensitivity analyses are used to calculate *R*_0_ and to determine which of the constituent parameters has the greatest influence on its magnitude.

## 2 Materials and methods

### 2.1 Basic reproduction number for LSDV

For an infection transmitted mechanically by biting insects the basic reproduction number can be written as,

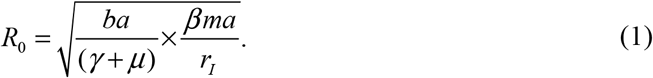

A full mathematical derivation of this expression for *R*_0_, including the transmission model on which it is based, is presented in the supporting information (Text S1). However, the expression, (1), can be understood heuristically as follows. After it feeds on an infected animal, an insect remains infected (and infectious) until the virus becomes inactivated or it dies, a period which lasts on average 1/(*γ*+*μ*) days, where *γ* is the virus inactivation rate and μ is the vector mortality rate. During this time it will bite susceptible cattle *a* times per day (where *a* is the reciprocal of the time interval between blood meals) and a proportion, *b*, of these bites (i.e. the probability of transmission from insect to bovine) will result in a newly infected host. Once infected, a bovine will remain infectious for the duration of its infectious period, which lasts 1/*r*_*I*_ days on average. During this time the host will be bitten by susceptible insects on average *m*×*a* times per day (here *m*=*N*/*H* is the vector to host ratio and *N* and *H* are the number of vectors and hosts, respectively), a proportion, *β*, of which will result in a newly infected vector (i.e. the probability of transmission from bovine to insect).

### 2.2 Mechanical transmission of LSDV by insects

Data on mechanical transmission of lumpy skin disease virus to cattle by five species of biting insect (*S. calcitrans*, *C. nubeculosus*, *Ae. aegypti*, *An. stephensi* and *Cx. quinquefasciatus*) were extracted from the published literature (Chihota, Rennie, Kitching & Mellor, 2001, 2003). This provided the number of positive insects (i.e. those for which viral DNA was detected; virus isolation was only carried out for a small number of pooled samples) and the number of insects tested after feeding on a LSDV-infected bovine (*S. calcitrans*, *C. nubeculosus* or *Ae. aegypti*) or a blood-virus mix via a membrane (*An. stephensi* and *Cx. quinquefasciatus*) at each day post feeding (Table S1). It also provided information on whether or not transmission occurred when batches of insects that had previously fed on an infected animal or received an infected blood meal were allowed to refeed on a naïve bovine (Table S1).

These data were used to estimate the virus inactivation rate (*γ*), the probability of transmission from bovine to insect (*β*) and the probability of transmission from insect to bovine (*b*) for each species. Parameters were estimated in a Bayesian framework to facilitate the incorporation of uncertainty in estimates of *R*_0_. The likelihood for the data is given by,

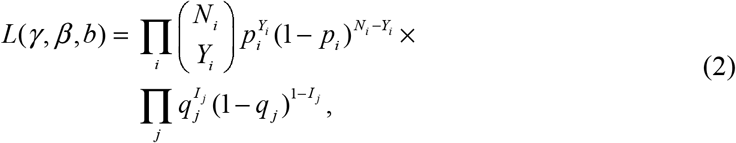

where *Y*_*i*_ and *N*_*i*_ are the number of positive insects and number of insects tested at *t*_*i*_ days post feeding, respectively, and *I*_*j*_ is a variable indicating whether (*I*_*j*_ =1) or not (*I*_*j*_=0) transmission occurred when insects were allowed to refeed on naïve animal *j* at *t*_*j*_ days post initial feed. In equation (2),

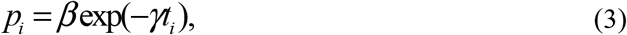

is the probability that an insect is positive at *t*_*i*_ days post feeding and,

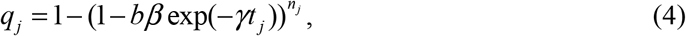

is the probability that transmission occurred from infected insects to bovine *j* at *t*_*j*_ days post initial feed. The probability, (3), is the probability that an insect became infected (*β*) multiplied by the probability that it was still infected when tested (exp(-*γt*_*i*_)). The probability, (4), is the probability that at least one insect (out of the *n*_*j*_ feeding) transmitted LSDV, where the probability that an individual insect will transmit LSDV is the product of the probabilities that it became infected (*β*), that it was still infected when refeeding occurred (exp(-*γt*_*j*_)) and that it subsequently transmitted LSDV to the animal during refeeding (*b*). Non-informative priors were assumed for all three parameters: exponential with mean 100 for *γ* and uniform with range (0,1) for *b* and *β*.

Samples from the joint posterior density were generated using an adaptive Metropolis scheme (Haario, Saksman & Tamminen, 2001), modified so that the scaling factor was tuned during burn-in to ensure an acceptance rate of between 20% and 40% for more efficient sampling of the target distribution (Andrieu & Thoms, 2008). Two chains of 50,000 iterations were run, with the preceding 10,000 iterations discarded to allow for burn-in of the chain. The chains were then thinned (taking every fifth sample) to reduce autocorrelation amongst the samples. The adaptive Metropolis scheme was implemented in Matlab (version R2018a; The Mathworks Inc.). Convergence of the scheme was assessed visually and by examining the Gelman-Rubin statistic provided in the coda package (Plummer, Best, Cowles & Vines, 2006) in R (R Core Team, 2018).

### 2.3 Latent and infectious periods for LSDV in cattle

The mean infectious period (1/*r*_*I*_) for LSDV in cattle was estimated using data from experimental infections (Tuppurainen, Venter & Coetzer, 2005; Babiuk et al., 2008). Three proxy measures of infectiousness were considered: detection of viral DNA in blood by PCR; detection of virus in blood by virus isolation (VI) in cell culture; and detection of virus (by transmission electron microscopy) or viral DNA (by PCR) in skin lesions. For each animal, the minimum infectious period was calculated as the time between the first positive and last positive samples, while the maximum infectious period was calculated as the time between the last negative and first subsequent negative sample for each measure (i.e. accounting for sampling frequency) (Table S2).

Although it is not needed for the calculation of *R*_0_ (see equation (1)), the mean latent period was also estimated for each proxy measure using data from experimental infections of cattle (Chihota, Rennie, Kitching & Mellor, 2001; Tuppurainen, Venter & Coetzer, 2005; Babiuk et al., 2008). In this case, the shortest latent period was the time of the last negative sample and the longest latent period was the time of the first positive sample for each measure (i.e. detection of viral DNA in blood, detection of virus in blood and appearance of skin lesions) (Table S2).

The infectious period was assumed to follow a gamma distribution with mean duration 1/*r*_*I*_ and shape parameter *n*_*I*_, while the latent period was assumed to follow a gamma distribution with mean duration 1/*r*_*E*_ and shape parameter *n*_*E*_ (see Text S1). Parameters (i.e. mean and shape parameter) for each proxy measure were estimated using Bayesian methods, with the likelihood given by,

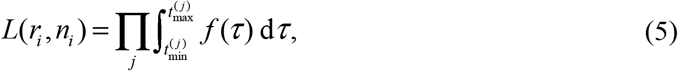

where *t*_min_ and *t*_max_ are the minimum and maximum infectious period (*i*=*I*) or latent period (*i*=*E*) for an animal, respectively, and *f* is the probability density function for the gamma distribution. Non-informative priors (exponential with mean 100) were assumed for both the mean and the shape parameter.

Samples from the joint posterior distribution were generated using a random walk Metropolis-Hastings algorithm (Andrieu & Thoms, 2008), with each parameter updated in turn. Two chains of 20,000 iterations were run, with the preceding 5,000 iterations discarded to allow for burn-in of the chain. The chains were then thinned (taking every other sample) to reduce autocorrelation amongst the samples. The Metropolis-Hastings scheme was implemented in Matlab (version R2018a; The Mathworks Inc.). Convergence of the scheme was assessed visually and by examining the Gelman-Rubin statistic provided in the coda package (Plummer, Best, Cowles & Vines, 2006) in R (R Core Team, 2018).

### 2.4 Vector life history parameters

Life history parameters, specifically the reciprocal of the time interval between blood meals (*a*), the vector to host ratio (*m*) and the vector mortality rate (μ), were estimated for *S. calcitrans*, *C. nubeculosus*, *Ae. aegypti*, *An. stephensi* and *Cx. quinquefasciatus*. For each parameter, plausible ranges were derived from the published literature (Table 1), so they could be incorporated in the uncertainty and sensitivity analysis.

**Table 1.** Life history parameters for five putative insect vectors of lumpy skin disease virus.

| parameter | symbol | range | comments | references |
| --- | --- | --- | --- | --- |
| *Stomoxys calcitrans* |  |  |  |  |
| biting rate (day<sup>-1</sup>) | *a* | 0.33-6 | time interval between blood meals (1/*a*) of 4-72 hours | Salem et al. (2013); Baldacchino et al. (2013) |
| vector to host ratio | *m* | 30-145 | - | Kunz & Monty (1976); Jacquet et al. (2014) |
| mortality rate (day<sup>-1</sup>) | $\mu$ | 0.04-0.11 | median lifespan (1/$\mu$) of 9-26 days | Salem et al. (2013); Skovgård & Nachman (2017) |
| *Culicoides nubeculosus* |  |  |  |  |
| biting rate (day<sup>-1</sup>) | *a* | 0-0.4 | based on duration of gonotrophic cycle | Birley & Boorman (1982); Braverman et al. (1985); Mullens et al. (2004) |
| vector to host ratio | *m* | 0-5000 | based on a maximum host biting rate ($m \times a$) of 2500 bites per host per day | Gerry et al. (2001) |
| mortality rate (day<sup>-1</sup>) | $\mu$ | 0.1-0.5 | based on proportion of females surviving gonotrophic cycle | Birley & Boorman (1982); Braverman et al. (1985) |
| *Aedes aegypti* |  |  |  |  |
| biting rate (day<sup>-1</sup>) | *a* | 0.18-0.23 | - | Scott et al. (2000); Liu-Helmersson et al. (2014) |
| vector to host ratio | *m* | 0-80 | modelled ratio of *Aedes* to cattle | Gachohi et al. (2016) |
| mortality rate (day<sup>-1</sup>) | $\mu$ | 0.07-0.2 | lifespan of 5-15 days estimated using a modelling analysis of 50 mark-release-recapture field studies | Brady et al. (2013) |
| *Anopheles stephensi* |  |  |  |  |
| biting rate (day<sup>-1</sup>) | *a* | 0.25-0.5 | time interval between blood meals (1/*a*) of 2-4 days | Killeen et al. (2000); Charlwood et al. (2016) |
| vector to host ratio | *m* | 0.6-60 | based on a host biting rate ($m \times a$) of around 2 to 120 bites per host per day | Killeen et al. (2000); Ryan et al. (2017) |
| mortality rate (day<sup>-1</sup>) | $\mu$ | 0.02-0.38 | daily survival probability ($\exp(-\mu)$) of 0.68-0.98 | Kiszewski et al. (2004) |
| *Culex quinquefasciatus* |  |  |  |  |
| biting rate (day<sup>-1</sup>) | *a* | 0.08-0.25 | time interval between blood meals<br>(1/*a*) of 4-12 days | Griffith & Turner (1996); Reisen et al. (2006) |
| vector to host ratio | *m* | 0-80 | modelled ratio of *Culex* to cattle | Gachohi et al. (2016) |
| mortality rate (day<sup>-1</sup>) | $\mu$ | 0.07-0.84 | - | Gad et al. (1989); Jones et al. (2012) |

### 2.5 Uncertainty and sensitivity analyses

Replicated Latin hypercube sampling (LHS) was used to explore the parameters influencing the basic reproduction number, *R*_0_ for each insect species and proxy measure of infectiousness (Blower & Dowlatabadi, 1994; Luz, Codeco, Massad & Struichner, 2003; Gubbins et al., 2008). Parameters were sampled either from their marginal posterior distributions (*b*, *β*, *γ* and 1/*r*_*I*_) or uniformly from plausible ranges (*a*, *m* and μ). The LHS results were used to compute the median and 95% prediction interval for *R*_0_. The sensitivity of *R*_0_ to changes in each parameter was assessed by calculating the partial rank correlation coefficients (PRCCs). The uncertainty and sensitivity analyses were implemented in Matlab (version R2018a; The Mathworks Inc.).

## 3 Results

### 3.1 Mechanical transmission of LSDV by insects

The model, (3), adequately captured the data for all five species of biting insect, with the observed number of positive insects lying within the 95% credible intervals for the posterior predictive distribution (Figure 1). Although there is uncertainty associated with all the parameters related to mechanical transmission, there are still clear differences amongst the species (Figure 2). The virus inactivation rate is lowest for *Ae. aegypti* (0.08 day^−1^), intermediate for *An. stephensi* (0.2 day^−1^) and *Cx. quinquefasciatus* (0.3 day^−1^) and highest for *S. calcitrans* (1.6 day^−1^) and *C. nubeculosus* (71.4 day^−1^) (Table 2). The probability of transmission from bovine to insect is highest for the three mosquito species, lower for *S. calcitrans* and lowest for *C. nubeculosus* (Table 2; Figure 2).

**Table 2.** Parameters for mechanical transmission of lumpy skin disease virus for five species of biting insect.

| species | virus inactivation rate<br>($\gamma$; day<sup>-1</sup>) | | mean duration of infection<br>(1/$\gamma$; days) | | probability of transmission<br>from bovine to insect ($\beta$) | | probability of transmission<br>from insect to bovine ($b$) | |
| --- | --- | --- | --- | --- | --- | --- | --- | --- |
|  | estimate† | 95% CI‡ | estimate | 95% CI | estimate | 95% CI | estimate | 95% CI |
| *S. calcitrans*§ | 1.71 | (0.80, 3.34) | 0.58 | (0.30, 1.26) | 0.46 | (0.23, 0.71) | 0.07 | (0.002, 0.64) |
| *C. nubeculosus*§ | 71.16 | (4.65, 390.0) | 0.01 | (0.003, 0.22) | 0.27 | (0.09, 0.54) | 0.50 | (0.03, 0.98) |
| *Ae. aegypti* | 0.09 | (0.01, 0.19) | 11.23 | (5.19, 79.11) | 0.90 | (0.64, 0.99) | 0.48 | (0.04, 0.97) |
| *An. stephensi* | 0.24 | (0.10, 0.42) | 4.08 | (2.38, 9.64) | 0.61 | (0.36, 0.83) | 0.03 | (0.001, 0.16) |
| *Cx. quinquefasciatus* | 0.34 | (0.17, 0.55) | 2.96 | (1.81, 5.74) | 0.72 | (0.47, 0.91) | 0.04 | (0.002, 0.28) |

† posterior median
‡ CI: credible interval
§ estimates are shown for the analysis assuming 100 insects refed when attempting transmission to cattle

**Figure 1.**
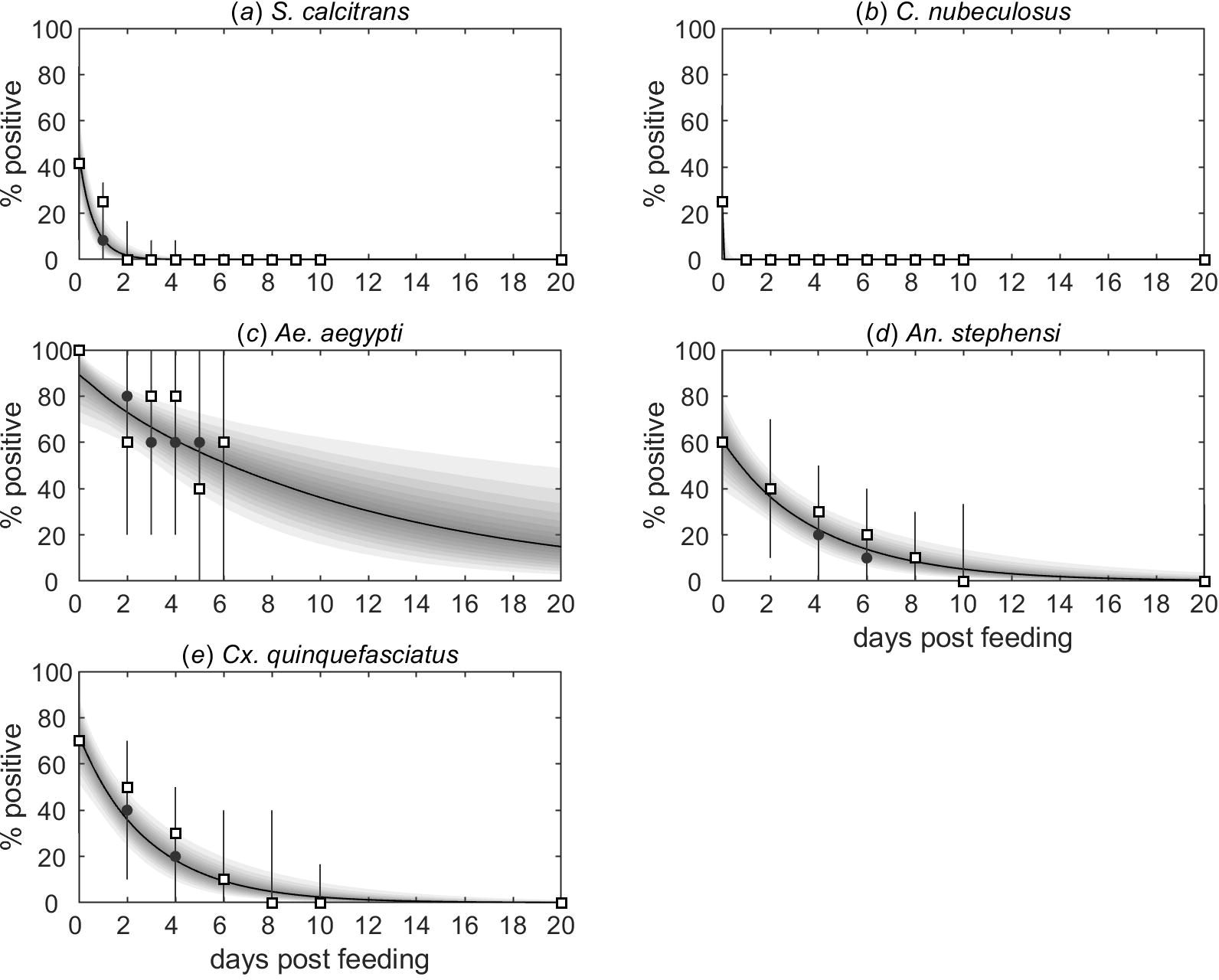
Proportion of biting insects positive for viral DNA after feeding on a bovine infected with lumpy skin disease virus. Results are shown for five species of biting insect: (*a*) *Stomoxys calcitrans*; (*b*) *Culicoides nubeculosus*; (*c*) *Aedes aegypti*; (*d*) *Anopheles stephensi*; and (*e*) *Culex quinquefasciatus*. Each plot shows the observed proportion of infected insects (open squares), the expected proportion of infected insects (black line: posterior median; shading: percentiles of the posterior distribution in 5% bands from 5% to 95%) and the posterior predictive distribution for the proportion of infected insects (grey circles: median; grey error bars: 95% prediction intervals).

**Figure 2.**
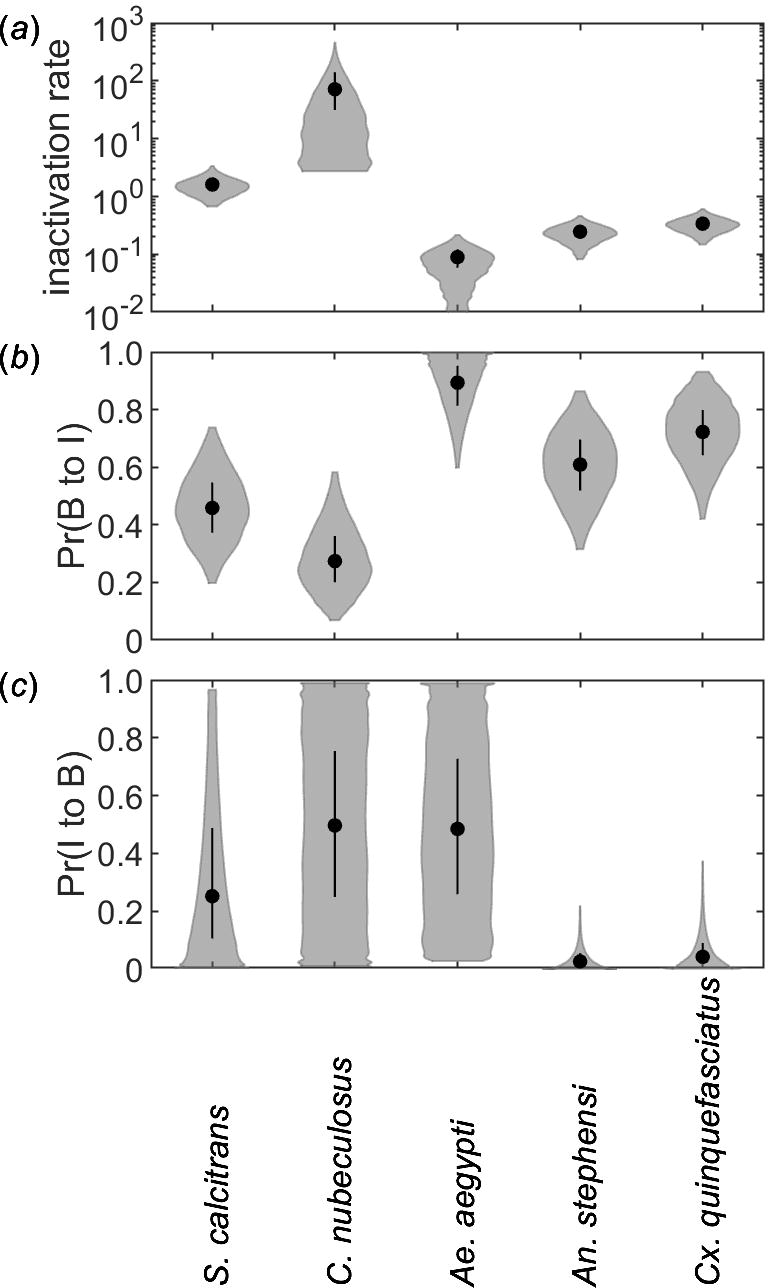
Parameters for mechanical transmission of lumpy skin disease virus by five species of biting insect: (*a*) virus inactivation rate (*γ*; day^−1^); (*b*) probability of transmission from bovine to insect (*β*; Pr(B to I)); and (*c*) probability of transmission from insect to bovine (*b*; Pr(I to B)). Each plot shows the median (black circle), 25th and 75th percentiles (black line) and density (shape) for the marginal posterior distribution. Results shown for *S. calcitrans* and *C. nubeculosus* assume 100 insects refed when attempting transmission to cattle.

There is considerable uncertainty in the estimates for the probability of transmission from insect to bovine for all five insect species (Figure 2; Table 2). The posterior mode was non-zero only for *Ae. aegypti*, though the probability of transmission from insect to bovine could not be precisely estimated for this species (Figure 2). The posterior median for the probability of transmission from insect to bovine was low for *An. stephensi* (0.03) and *Cx. quinquefasciatus* (0.04), though the upper 95% credible limit for both species is an order of magnitude higher (Figure 2; Table 2). Finally, the probability of transmission from insect to bovine could not be reliably estimated for either *S. calcitrans* or *C. nubeculosus* (Figure 2; Figure S1). This is, in part, because the number of insects refeeding when attempting transmission to cattle was not reported for these species (Chihota, Rennie, Kitching & Mellor, 2003). For *C. nubeculosus*, the posterior distribution for the probability of transmission from insect to bovine was identical to the prior distribution (Figure S1), regardless of assumptions made about the numbers of insects that refed (either 1, 5, 10, 20, 50 or 100). For *S. calcitrans* the posterior mass for the probability of transmission from insect to bovine was shifted towards zero if a larger number of insects was assumed to refeed, though the 95% credible interval remained large (Figure S1). For both species, estimates for the virus inactivation rate and probability of transmission from bovine to insect were not affected by the number of insects assumed to refeed (Figure S1). In the uncertainty and sensitivity analyses for these species, the posterior distributions for parameters related to mechanical transmission (*b*, *β* and *γ*) were those obtained when 100 insects were assumed to refeed, as this provides the most conservative assessment of the risk they pose.

### 3.2 Latent and infectious periods for LSDV in cattle

The mean infectious period depends on the proxy measure used to determine when an animal is infectious (Table 3). It is shortest when based on virus isolation from blood (8.8 days), intermediate when based on detection of viral DNA in blood (16.3 days) and longest when based on detection of virus or viral DNA in skin lesions (23.1 days). The mean latent period also depends on the proxy measure used to determine when an animal is infectious. It was estimated to be 5.8 days for blood (PCR), 8.1 days for blood (VI) and 7.3 days for skin lesions (Table 3).

**Table 3.**
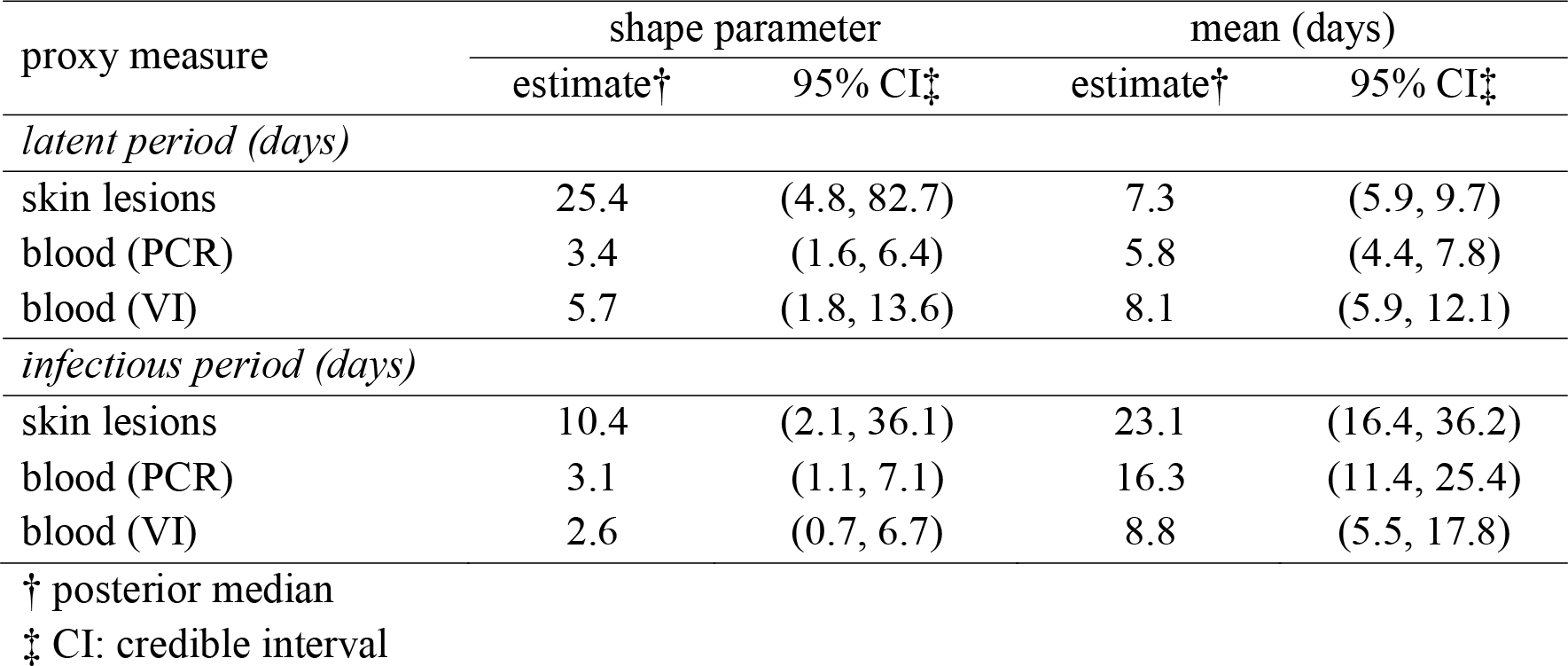
Parameters for latent and infectious periods for lumpy skin disease virus in cattle.

| proxy measure | shape parameter |  | mean (days) |  |
| --- | --- | --- | --- | --- |
|  | estimate<sup>†</sup> | 95% CI<sup>‡</sup> | estimate<sup>†</sup> | 95% CI<sup>‡</sup> |
| *latent period (days)* |  |  |  |  |
| skin lesions | 25.4 | (4.8, 82.7) | 7.3 | (5.9, 9.7) |
| blood (PCR) | 3.4 | (1.6, 6.4) | 5.8 | (4.4, 7.8) |
| blood (VI) | 5.7 | (1.8, 13.6) | 8.1 | (5.9, 12.1) |
| *infectious period (days)* |  |  |  |  |
| skin lesions | 10.4 | (2.1, 36.1) | 23.1 | (16.4, 36.2) |
| blood (PCR) | 3.1 | (1.1, 7.1) | 16.3 | (11.4, 25.4) |
| blood (VI) | 2.6 | (0.7, 6.7) | 8.8 | (5.5, 17.8) |

<sup>†</sup> posterior median
<sup>‡</sup> CI: credible interval

### 3.3 Uncertainty and sensitivity analyses

The basic reproduction number for LSDV was highest for transmission by *S. calcitrans*, intermediate for *Ae. aegypti*, low for *C. nubeculosus* and *An. stephensi* and lowest *Cx. quinquefasciatus* (Figure 3; Table 4). Indeed, the median *R*_0_ for *Cx. quinquefasciatus* was below the threshold for an outbreak to occur (i.e. *R*_0_=1). The magnitude of *R*_0_ depended on the proxy measure of infectiousness used (through its influence on the duration of infectiousness), with higher values when based on detection of virus or viral DNA in skin lesions, intermediate when based on detection of viral DNA in blood and lowest when based on detection of virus in blood (Figure 3; Table 4).

**Table 4.**
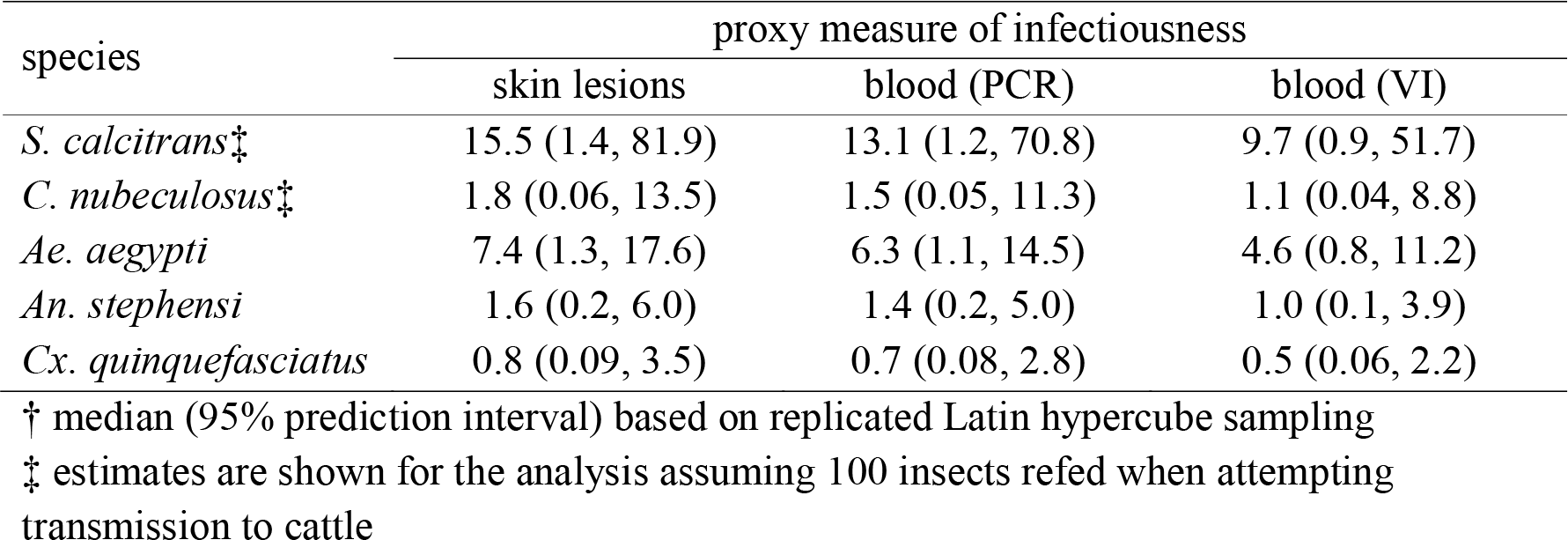
Basic reproduction number† (*R*_0_) for lumpy skin disease virus (LSDV) for five species of biting insect.

**Figure 3.**
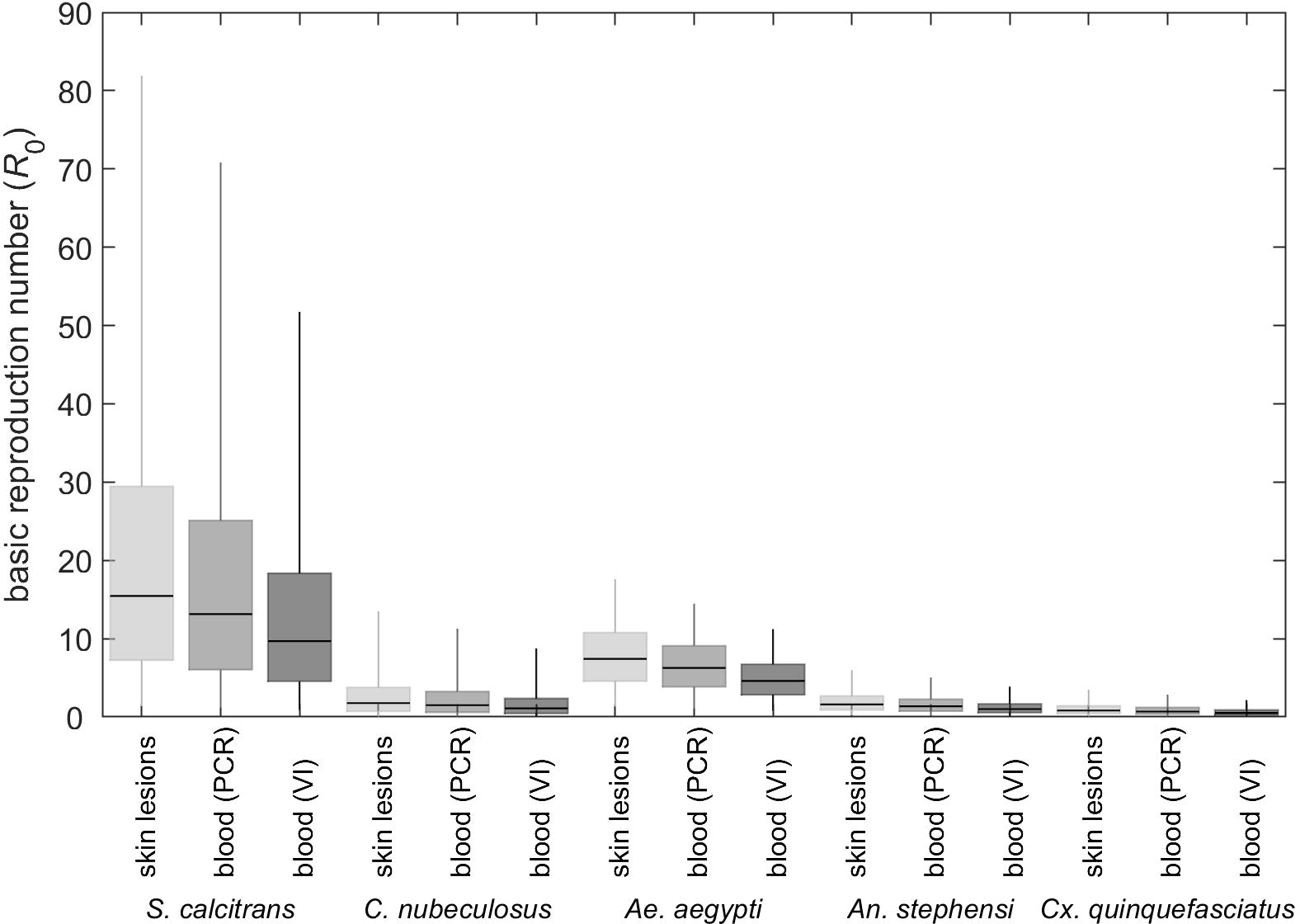
Basic reproduction number (*R*_0_) for lumpy skin disease virus (LSDV) when transmitted by *Stomoxys calcitrans*, *Culicoides nubeculosus*, *Aedes aegypti*, *Anopheles stephensi* or *Culex quinquefasciatus*. The estimated values for *R*_0_ are shown for three proxy measures of infectiousness: detection of virus or viral DNA in skin lesions (light grey); detection of viral DNA in blood (mid grey); or detection of virus in blood (dark grey). Box and whisker plots show the median (black horizontal line), interquartile range (coloured box) and 95% range (whiskers) based on replicated Latin hypercube sampling (100 replicates with the range for each parameter subdivided into 100 steps). Results shown for *S. calcitrans* and *C. nubeculosus* used the posterior distributions assuming 100 insects refed when attempting transmission to cattle.

The sensitivity of *R*_0_ to changes in its constituent parameters differed amongst the insect species, though some patterns emerge (Figure 4). For all species, the strongest correlations were with the probability of transmission from insect to bovine (*b*; PRCC>0.66), the biting rate (*a*; PRCC>0.75, except for *Ae. aegypti*), the vector to host ratio (*m*; PRCC>0.58) and the virus inactivation rate (*γ*; PRCC<−0.47, except *An. stephensi* and *Cx. quinquefasciatus*). For the three mosquito species, there was also correlation with the vector mortality rate (μ; PRCC<-0.45), but this was not the case for the other dipteran species (PRCC≈0). The probability of transmission from bovine to insect (*β*) and the mean infectious period (1/*r*_*I*_) were also positively correlated with *R*_0_ (PRCC=0.3-0.4 and PRCC=0.2-0.5, respectively). These patterns are independent of the proxy measure of infectiousness used (Figure 4).

**Figure 4.**
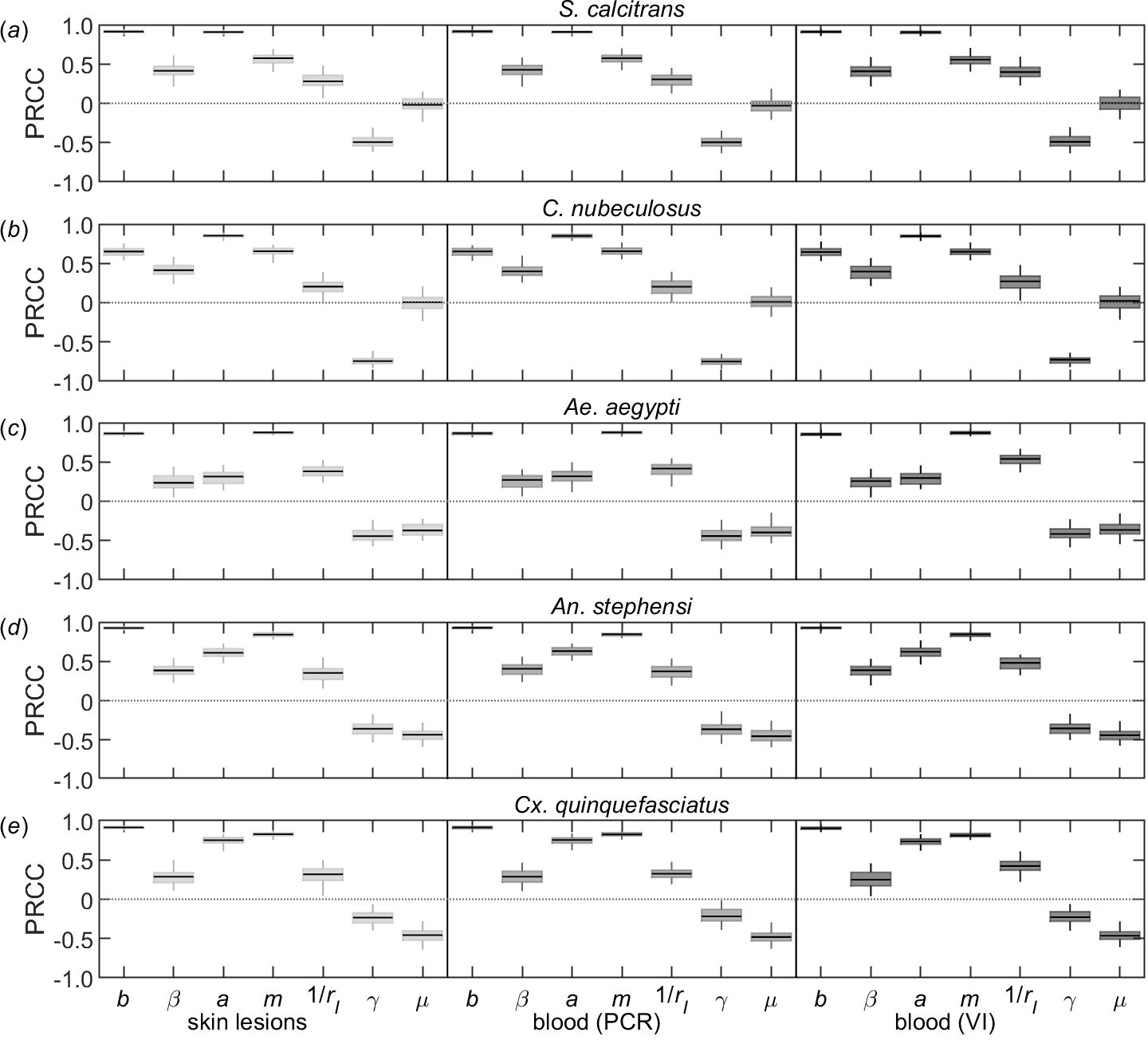
Sensitivity analysis of the basic reproduction number (*R*_0_) for lumpy skin disease virus (LSDV) when transmitted by: (*a*) *Stomoxys calcitrans*; (*b*) *Culicoides nubeculosus*; (*c*) *Aedes aegypti*; (*d*) *Anopheles stephensi*; or (*e*) *Culex quinquefasciatus*. Plots show the partial rank correlation coefficients (PRCC) for each parameter when the estimated values for *R*_0_ are calculated using three proxy measures of infectiousness: detection of virus or viral DNA in skin lesions (light grey); detection of viral DNA in blood (mid grey); or detection of virus in blood (dark grey). Box and whisker plots show the median (black horizontal line), interquartile range (coloured box) and 95% range (whiskers) for the PRCCs based on replicated Latin hypercube sampling (100 replicates with the range for each parameter subdivided into 100 steps). Results shown for *S. calcitrans* and *C. nubeculosus* used the posterior distributions assuming 100 insects refed when attempting transmission to cattle.

## 4 Discussion

The results of the uncertainty analysis for the basic reproduction number (Figure 3; Table 4) suggest that *S. calcitrans* is likely to be the most efficient vector of LSDV (out of the five species considered in the present study), with *Ae. aegypti* also an efficient vector of the virus. By contrast, *C. nubeculosus*, *An. stephensi* and *Cx. quinquefasciatus* are likely to be inefficient as vectors of LSDV.

These conclusions would potentially be different if the risk posed by each species was assessed on the outcome of the transmission experiments alone (Table 2). While the experimental outcome suggests *Ae. aegypti* might be an efficient vector, it could be concluded that *S. calcitrans* is likely to be a poor vector given the very short duration of infection (<1 day) and the comparatively low probabilities of transmission from bovine to insect or insect to bovine for this species. By contrast, the results of the transmission experiments suggest that *Cx. quinquefasciatus* could be a more efficient vector than it is based on the estimates for *R*_0_, because of the relatively long duration of infection and the high probability of transmission from bovine to insect. This highlights the importance of considering all factors associated with transmission, including vector life history, when assessing the risk of transmission for each species. For both *S. calcitrans* and *Cx. quinquefasciatus* the main reason for the difference in conclusions between transmission experiments and *R*_0_ is the time interval between blood meals, which is shorter than the duration of infection for *S. calcitrans*, but longer than the duration of infection for *Cx. quinquefasciatus* (Tables 1 & 2).

A certain amount of care should be taken in interpreting these results, however, because there is substantial uncertainty associated with the estimates of *R*_0_ for all five species, but especially for *S. calcitrans* and *C. nubeculosus*. One of the major sources of uncertainty is the probability of transmission from insect to bovine (Figure 2; Table 2). Although transmission to naïve cattle was attempted (Chihota, Rennie, Kitching & Mellor, 2003), it was done at times when *S. calcitrans* and *C. nubeculosus* were unlikely to still be infectious (see Figure 1; refeeding was at 1-3 days post infection for *S. calcitrans* and 3-5 days post infection for *C. nubeculosus*). Consequently, there is little information in the data to estimate this parameter, which is reflected in the posterior distributions being almost the same as the prior distributions, especially in the case of *C. nubeculosus* (Figure 2; Figure S1). Yet the probability of transmission from insect to bovine was identified as one of the parameters to which the basic reproduction number is most sensitive for all five vector species considered (Figure 4).

The influence of the virus inactivation rate and vector mortality rate on *R*_0_ differed amongst the species (Figure 4). The duration of infection (i.e. the reciprocal of the virus inactivation rate) for *C. nubeculosus* or *S. calcitrans* is much shorter than the lifespan of an insect and, consequently, only the virus inactivation rate is important for determining the magnitude of *R*_0_. By contrast, the duration of infection and insect lifespan are comparable for *Ae. aegypti*, *An. stephensi* and *Cx. quinquefasciatus*, so that both of these parameters have an influence on the basic reproduction number.

As with all uncertainty and sensitivity analyses, the conclusions drawn from them are valid only over the parameter ranges considered. For parameters related to mechanical transmission, these ranges were estimated from the available data on the outcome of transmission experiments (Chihota, Rennie, Kitching & Mellor, 2001, 2003) and so represent the best current estimates. Given the uncertainty in these parameters (which reflects the small numbers of animals and insects tested at each time point in the studies; Table S1), a focus of future experimental work should be to more precisely measure parameters related to mechanical transmission and, in particular, the probability of transmission from insect to bovine and the virus inactivation rate.

All three vector life history parameters were identified as influencing the magnitude of *R*_0_ (Figure 4). Plausible ranges for these parameters were drawn from the published literature (Table 1). Wherever possible, ranges were derived using data relating to the species themselves. If suitable data were not available, however, ranges were derived using data for related species. In addition, the vector life parameters were assumed to be constant, but they will depend on environmental factors, including temperature and rainfall. Consequently, the basic reproduction number, and so the risk of transmission, is unlikely to be constant over space or time, but will vary both geographically and seasonally.

This study considered the potential of five species of biting insects to transmit LSDV. This was primarily because their ability to transmit LSDV had been assessed experimentally for each of the species. However, the present analysis did not consider the effect of vector feeding preferences on their potential role in transmission. The stable fly, *S. calcitrans*, is one of the most damaging arthropod pests of cattle worldwide, both through its impact on production (Taylor, Moon & Mark, 2012) and its ability to transmit a number of pathogens (Baldacchino et al., 2013). Consequently, this species’ feeding preferences are unlikely to affect its efficiency as a vector of LSDV. Similarly, *C. nubeculosus* and other *Culicoides* species for which *C. nubeculosus* can be considered a model are livestock-associated (Lassen, Nielsen & Kristensen, 2012; Purse, Carpenter, Venter, Bellis & Mullens, 2015), suggesting host preference will not reduce their efficiency as a vector of LSDV. By contrast, *Ae. aegypti* feeds primarily on humans (Reiter, 2010), potentially limiting its role in the transmission of LSDV, despite its apparent efficiency as a vector. However, there are other *Aedes* species (for example, *Ae. cinereus* or *Ae. vexans*), which display a preference for feeding on cattle (Hayes, Tempelis, Hess & Reeves, 1973; Magnarelli, 1977) and for which *Ae. aegypti* could act as a model vector species. Many *Anopheles* species, including those in the Maculipennis subgroup, which is of most relevance to Europe (Sinka et al., 2010; ECDC, 2019), are described as zoophilic (Sinka et al., 2010). Accordingly, host preference is unlikely to limit their ability to transmit LSDV. Finally, host preference in *Cx. quinquefasciatus* varies according to environmental conditions and geographical area, though it does feed on mammals (Hayes, Tempelis, Hess & Reeves, 1973; Molaei, Huang & Andreadis, 2012; Janssen et al., 2015). However, any preference for non-bovine hosts in this species (or other *Culex* species for which it is a suitable model) will only further reduce its importance in the transmission of LSDV.

The magnitude of the basic reproduction number depends on the proxy measure used for infectiousness (Figure 3). Viral titres in blood can be low and detection of virus intermittent (Tuppurainen, Venter & Coetzer, 2005; Babiuk et al., 2008), suggesting this may not be the best proxy measure. By contrast, skin nodules have high titres of virus (Babiuk et al., 2008) and insects which feed on nodules become infected (Chihota, Rennie, Kitching & Mellor, 2001, 2003), indicating that this may be a reasonable proxy measure. This could be tested experimentally by feeding insects on areas with and without lesions of clinically-affected cattle and on infected, but subclinical cattle and relating viral titres in skin and blood to the probability of insects becoming infected.

The expression for the basic reproduction number presented in equation (1) assumes there is negligible disease-associated mortality in cattle (see Text S1). Reported mortality for LSD outbreaks in the Balkans was very low, with the median within-herd mortality being zero in all affected countries (EFSA, 2017). In endemic settings, mortality is generally low (1-3%) (Tuppurainen & Oura, 2012; Molla, Frankena & de Jong, 2017), but it may sometimes reach 40% (Tuppurainen & Oura, 2012). If disease-associated mortality (up to 40%) is included in the computation of *R*_0_, it does not greatly affect the conclusions of the present study regarding either the magnitude of *R*_0_ for each vector species or the sensitivity of *R*_0_ to changes in its constituent parameters (see Text S2).

Two previous studies have estimated the basic reproduction number for LSDV using outbreak data. In the first, Magori-Cohen et al. (2012)^1^ estimated *R*_0_ to be around 4 for an outbreak in a large dairy herd in Israel in 2006, where *S. calcitrans* was likely to be the principal vector species (Kahana-Sutin, Klement, Lensky & Gottlieb, 2017). Because of the control measures implemented on the farm (removal of cattle showing generalised disease on the day of detection), this implicitly assumed an infectious period of around one day. If a mean infectious period of 23 days is used instead (i.e. the maximum estimated in the present study; Table 3), the corresponding value for *R*_0_ is around 19, which is similar to the median estimate for transmission by *S. calcitrans* obtained in the present study (Figure 3; Table 4). By contrast, the second study estimated *R*_0_ to be around 1.1 for outbreaks in eight cattle herds in Ethiopia in 2014-2015, using viraemia as a proxy for infectiousness (Molla, Frankena & de Jong, 2017). This is substantially lower than the ranges estimated for either *S. calcitrans* or *Ae. aegypti*, but is consistent with those for *C. nubeculosus*, *An. stephensi* or *Cx. quinquefasciatus* (Figure 3; Table 4). However, the vector species involved in transmission of LSDV in Ethiopia are not known (Molla, de Jong & Frankena, 2017).

In this paper, the risk of transmission of LSDV was assessed for five species of biting insect using the basic reproduction number. The results suggest that *S. calcitrans* and *Ae. aegypti* are likely to be efficient vectors (i.e. *R*_0_ is substantially above one), while *C. nubeculosus*, *An. stephensi* and *Cx. quinquefasciatus* are likely to be inefficient vectors (i.e. *R*_0_ is close to or below one). However, there is considerable uncertainty associated with the estimates of *R*_0_ for LSDV and future work should focus in particular on estimating the probability of transmission from insect to bovine and the virus inactivation rate. Finally, using the basic reproduction number has demonstrated that any assessment of the risk posed by an insect vector needs to consider all factors which influence its ability to transmit the virus, including life history parameters.

## Acknowledgements

The author is grateful to Pip Beard, Beatriz Sanz-Bernardo and Christopher Sanders (all at The Pirbright Institute) for helpful comments on the manuscript.

1 Because of differences in model formulation, the value of *R*_0_ presented by Magori-Cohen et al. (2012) is the square of *R*_0_ in the current paper. Here the square root transformed values are presented.

## References

Alkhamis, M.A. & VanderWaal, K. (2016). Spatial and temporal epidemiology of lumpy skin disease in the Middle East, 2012-2015. Frontiers in Veterinary Science, 3, 19. https://doi.org/10.3389/fvets.2016.00019

Andrieu, C. & Thoms, J. (2008). A tutorial on adaptive MCMC. Statistics and Computing, 18, 343–373. https://doi.org/10.1007/s11222-008-9110-y

Babiuk, S., Bowden, T.R., Parkyn. G., Dalman, B., Manning, L., Neufeld, J., Embury-Hyatt, C., Copps, J. & Boyle, D.B. (2008). Quantification of lumpy skin disease virus following experimental infection in cattle. Transboundary and Emerging Diseases, 55, 299–307. https://doi.org/10.1111/j.1865-1682.2008.01024.x

Baldacchino, F., Muenwom, V., Desquesnes, M., Desoli, F., Charoenviriyaphap, T. & Duvallet, G. (2013). Transmission of pathogens by *Stomoxys* flies (Diptera: Muscidae): a review. Parasite, 20, 26. https://doi.org/10.1051/parasite/2013026

Birley, M.H. & Boorman, J.P.T. (1982). Estimating the survival and biting rates of hematophagous insects, with particular reference to the *Culicoides obsoletus* group (Diptera, Ceratopogonidae) in southern England. Journal of Animal Ecology, 51, 135–148. https://doi.org/10.2307/4315

Blower, S.M. & Dowlatabadi, H. (1994). Sensitivity and uncertainty analysis of complex models of disease transmission: an HIV model as an example. International Statistical Review, 62, 229–243. https://doi.org/10.2307/1403510

Brady, O.J., Johansson, M.A., Guerra, C.A., Bhatt, S., Golding, N., Pigott, D.M., Delatte, H., Grech, M.G., Leisnham, P.T., Maciel-de-Freitas, R., Styer, L.M., Smith, D.L., Scott, T.W., Gething, P.W. & Hay, S.I. (2013). Modelling adult *Aedes aegypti* and *Aedes albopictus* survival at different temperatures in laboratory and field settings. Parasites & Vectors, 6, 351. https://doi.org/10.1186/1756-3305-6-351

Braverman, Y., Linley, J. R., Marcus, R. & Frish, K. (1985). Seasonal survival and expectation of infectious life of *Culicoides* spp. (Diptera: Ceratopogonidae) in Israel, with implications for bluetongue virus transmission and a comparison of the parous rate in *C. imicola* from Israel and Zimbabwe. Journal of Medical Entomology, 22, 476–484. https://doi.org/10.1093/jmedent/22.5.476

Carn, V.M. & Kitching, R.P. (1995). An investigation of possible routes of transmission of lumpy skin disease virus (Neethling). Epidemiology and Infection, 114, 219–226. https://doi.org/10.1017/S0950268800052067

Charlwood, J.D., Nenhep, S., Sovannaroth, S., Morgan, J.C., Hemingway, J., Chitnis, N. & Briët, J.T. (2016). ‘Nature or nurture’: survival rate, oviposition interval and possible gonotrophic discordance among South East Asian anophelines. Malaria Journal, 15, 356. https://doi.org/10.1186/s12936-016-1389-0

Chihota, C.M., Rennie, L.F., Kitching, R.P. & Mellor, P.S. (2001). Mechanical transmission of lumpy skin disease virus by *Aedes aegypti* (Diptera: Culicidae). Epidemiology and Infection, 126, 317–321. https://doi.org/10.1017/S0950268801005179

Chihota, C.M., Rennie, L.F., Kitching, R.P. & Mellor, P.S. (2003). Attempted mechanical transmission of lumpy skin disease virus by biting insects. Medical and Veterinary Entomology, 17, 294–300. https://doi.org/10.1046/j.1365-2915.2003.00445.x

Diekmann, O. & Heesterbeek, J.A.P. (2000). Mathematical epidemiology of infectious diseases: model building, analysis and interpretation. Chichester: John Wiley & Sons.

ECDC (2019). European Centre for Disease Prevention and Control and European Food Safety Authority: Mosquito maps. https://ecdc.europa.eu/en/disease-vectors/surveillance-and-disease-data/mosquito-maps

EFSA (2017). Scientific report on lumpy skin disease: I. Data collection and analysis. The EFSA Journal, 15, 4773. https://doi.org/10.2903/j.efsa.2017.4773

EFSA Panel on Animal Health and Welfare (2015). Scientific opinion on lumpy skin disease. The EFSA Journal, 13, 3986. https://doi.org/10.2903/j.efsa.2015.3986

Gachohi, J.M., Njenga, M.K., Kitala, P. & Bett, B. (2016). Modelling vaccinations strategies against Rift Valley fever virus in livestock in Kenya. PLoS Neglected Tropical Diseases, 10, e0005049. https://doi.org/10.1371/journal.pntd.0005049

Gad, A.M., Feinsod, F.M., Soliman, B.A. & El Said, S. (1989). Survival estimates for adult *Culex pipiens* in the Nile Delta. Acta Tropica, 46, 173–179. https://doi.org/10.1016/0001-706X(89)90034-X

Gerry, A.C., Mullens, B.A., MacLachlan, N.J. & Mecham, O.J. (2001). Seasonal transmission of bluetongue virus by *Culicoides sonorensis* (Diptera: Ceratopogonidae) at a southern California dairy and evaluation of vectorial capacity as a predictor of bluetongue virus transmission. Journal of Medical Entomology, 38, 197–209. https://doi.org/10.1603/0022-2585-38.2.197

Griffith, J.S.R. & Turner, G.D. (1996). Culturing *Culex quinquefasciatus* mosquitoes with a blood substitute diet for the females. Medical and Veterinary Entomology, 10, 265–268. https://doi.org/10.1111/j.1365-2915.1996.tb00741.x

Gubbins, S., Carpenter, S., Baylis, M., Wood, J.L.N. & Mellor, P.S. (2008). Assessing the risk of bluetongue to UK livestock: uncertainty and sensitivity analysis of a temperature-dependent model for the basic reproduction number. Journal of the Royal Society Interface, 5, 363–371. https://doi.org/10.1098/rsif.2007.1110

Haario, H., Saksman, E. & Tamminen, J. (2001). An adaptive Metropolis algorithm. Bernoulli, 7, 223–242. https://projecteuclid.org/euclid.bj/1080222083

Hayes, R.O., Tempelis, C.H., Hess, A.D. & Reeves, W.C. (1973). Mosquito preference studies in Hale County, Texas. American Journal of Tropical Medicine and Hygiene, 22, 270–277. https://doi.org/10.4269/ajtmh.1973.22.270

Jacquiet, P., Rouet, D., Bouhsira, E., Salem, A., Lienard, E. & Franc, M. (2014). Population dynamics of *Stomoxys calcitrans* (L.) (Diptera: Muscidae) in southwestern France. Revue de Médicine Vétérinaire, 165, 267–271. https://www.revmedvet.com/artdes-us.php?id=16015

Janssen, N., Fernandez-Salas, I., Díaz González, E.E., Gaytan-Burns, A., Medina-de la Garza, C.E., Sanchez-Casas, R.M., Börstler, J., Cadar, D., Schmidt-Chanasit, J. & Jöst, H. (2015). Mammalophilic feeding behaviour of *Culex quinquefasciatus* mosquitoes collected in the cities of Chetumal and Cancun, Yucatán Peninsula, Mexico. Tropical Medicine and International Health, 20, 1488–1491. https://doi.org/10.1111/tmi.12587

Jones, C.E., Lounibos, P., Marra, P.P. & Kilpatrick, A.M. (2012). Rainfall influences survival of *Culex pipiens* (Diptera: Culicidae) in a residential neighbourhood in the mid-Atlantic United States. Journal of Medical Entomology, 49, 467–473. https://doi.org/10.1603/ME11191

Kahana-Sutin, E., Klement, E., Lensky, I. & Gottlieb, Y. (2017). High relative abundance of the stable fly *Stomoxys calcitrans* is associated with lumpy skin disease outbreaks in Israeli dairy farms. Medical and Veterinary Entomology, 31, 150–160. https://doi.org/10.1111/mve.12217

Killeen, G.F., McKenzie, F.E., Foy, B.D., Shieffelin, C., Billingsley, P.F. & Beier, J.C. (2000). A simplified model for predicting malaria entomologic inoculation rates based on entomologic and parasitologic parameters relevant to control. American Journal of Tropical Medicine and Hygiene, 62, 535–544. https://doi.org/10.4269/ajtmh.2000.62.535

Kiszewski, A., Mellinger, A., Spielman, A., Malaney, P., Ehrlich Sachs, S. & Sachs, J. (2004). A global index representing the stability of malaria transmission. American Journal of Tropical Medicine and Hygiene, 70, 486–498. https://doi.org/10.4269/ajtmh.2004.70.486

Kitching, R.P. & Mellor, P.S. (1986). Insect transmission of capripox virus. Research in Veterinary Science, 40, 255–258. https://doi.org/10.1016/S0034-5288(18)30523-X

Kunz, S.E. & Monty, J. (1976). Biology and ecology of *Stomoxys nigra* Macquart and *Stomoxys calcitrans* (L.) (Diptera, Muscidae) in Mauritius. Bulletin of Entomological Research, 66, 745–755. https://doi.org/10.1017/S0007485300010798

Lassen, S.B., Nielson, S.A. & Kristensen, M. (2012). Identity and diversity of blood meal hosts of biting midges (Diptera: Ceratopogonidae: *Culicoides* Latreille) in Denmark. Parasites & Vectors, 5, 143. https://doi.org/10.1186/1756-3305-5-143

Liu-Helmersson, J., Stenlund, H., Wilder-Smith, A. & Rocklöv, J. (2014). Vectorial capacity of *Aedes aegypti*: effects of temperature and implications for global dengue epidemic potential. PLoS ONE, 9, e89783. https://doi.org/10.1371/journal.pone.0089783

Luz, P.M., Codeco, C.T., Massad, E. & Struichner, C.J. (2003). Uncertainties regarding dengue modelling in Rio de Janeiro, Brazil. Memorias do Instituto Oswaldo Cruz, 98, 871–878. https://doi.org/10.1590/S0074-02762003000700002

Magnarelli, L.A. (1977). Host feeding patterns of Connecticut mosquitoes (Diptera: Culicidae). American Journal of Tropical Medicine and Hygiene, 26, 547–552. https://doi.org/10.4269/ajtmh.1977.26.547

Magori-Cohen, R., Louzoun, Y., Herziger, Y., Oron, E., Arazi, A., Tuppurainen, E., Shpigel, N.Y. & Klement, E. (2012). Mathematical modelling and evaluation of the different routes of transmission of lumpy skin disease virus. Veterinary Research, 43, 1. https://doi.org/10.1186/1297-9716-43-1

Mellor, P.S., Kitching, R.P. & Wilkinson, P.J. (1987). Mechanical transmission of capripox virus and African swine fever virus by *Stomoxys calcitrans*. Research in Veterinary Science, 43, 109–112. https://doi.org/10.1016/S0034-5288(18)30753-7

Mercier, A., Arsevska, E., Bournez, L., Bronner, A., Calavas, D., Cauchard, J., Falala, S., Caufour, P., Tisseuil, C., Lefrançois, T. & Lancelot, R. (2018). Spread rate of lumpy skin disease in the Balkans, 2015-2016. Transboundary and Emerging Diseases, 65, 240–243. https://doi.org/10.1111/tbed.12624

Molaei, G., Huang, S. & Andreadis, T.G. (2012). Vector-host interactions of *Culex pipiens* complex in northeastern and southwestern USA. Journal of the American Mosquito Control Association, 28(Suppl4), 127–136. https://doi.org/10.2987/8756-971X-28.4s.127

Molla, W., de Jong, M.C.M. & Frankena, K. (2017). Temporal and spatial distribution of lumpy skin disease outbreaks in Ethiopia in the period 2000 to 2015. BMC Veterinary Research, 13, 310. https://doi.org/10.1186/s12917-017-1247-5

Molla, W., Frankena, K. & de Jong, M.C.M. (2017). Transmission dynamics of lumpy skin disease in Ethiopia. Epidemiology and Infection, 145, 2856–2863. https://doi.org/10.1017/S0950268817001637

Molla, W., de Jong, M.C.M., Gari, G. & Frankena, K. (2017). Economic impact of lumpy skin disease and cost effectiveness of vaccination for the control of outbreaks in Ethiopia. Preventive Veterinary Medicine, 147, 100–107. https://doi.org/10.1016/j.prevetmed.2017.09.003

Mullens, B.A., Gerry, A.C., Lysyk, T.J. & Schmidtmann, E.T. (2004). Environmental effects on vector competence and virogenesis of bluetongue virus in *Culicoides*: interpreting laboratory data in a field context. Veterinaria Italiana, 40, 160–166. http://www.izs.it/vet_italiana/2004/40_3/32.pdf

Plummer, M., Best, N., Cowles, K. & Vines, K. (2006). CODA: Convergence Diagnosis and Output Analysis for MCMC. R News, 6, 7–11.https://cran.r-project.org/doc/Rnews/Rnews_2006-1.pdf

Purse, B.V., Carpenter, S., Venter, G.J., Bellis, G. & Mullens, B.A. (2015). Bionomics of temperate and tropical *Culicoides* midges: knowledge gaps and consequences for transmission of Culicoides-borne viruses. Annual Review of Entomology, 60, 373–392. https://doi.org/10.1146/annurev-ento-010814-020614

R Core Team 2018 R: A language and environment for statistical computing. R Foundation for Statistical Computing, Vienna, Austria. http://www.R-project.org/

Reisen, W.K., Fang, Y. & Martinez, V.M. (2006). Effects of temperature on the transmission of West Nile virus by *Culex tarsalis* (Diptera: Culicidae). Journal of Medical Entomology, 43, 309–317. https://doi.org/10.1093/jmedent/43.2.309

Reiter, P. (2010). Yellow fever and dengue: a threat to Europe? EuroSurveillance, 15, 19509. https://www.eurosurveillance.org/content/10.2807/ese.15.10.19509-en

Ryan, S.J., Lippi, C.A., Boersch-Supan, P.H., Heydari, N., Silva, M., Adrian, J., Noblecilla, L.F., Ayala, E.B., Encalada, M.D., Larsen, D.A., Krisher, J.T., Krisher, L., Fregosi, L. & Stewart-Ibarra, A.M. (2017). Quantifying seasonal and diel variation in Anopheline and Culex human biting rates in Southern Ecuador. Malaria Journal, 16, 479. https://doi.org/10.1186/s12936-017-2121-4

Salem, A., Franc, M., Jacquiet, P., Bouhsira, E. & Lienard, E. (2012). Feeding and breeding aspects of *Stomoxys calcitrans* (Diptera: Muscidae) under laboratory conditions. Parasite, 19, 309–317. https://doi.org/10.1051/parasite/2012194309

Scott, T.W., Amerasinghe, P.H., Morrison, A.C., Lorenz, L.H., Clark, G.G., Strickman, D., Kittayapog, P. & Edman, J.D. (2000). Longitudinal studies of *Aedes aegypti* (Diptera: Culicidae) in Thailand and Puerto Rico: blood feeding frequency. Journal of Medical Entomology, 37, 89–101. https://doi.org/10.1603/0022-2585-37.1.77

Sinka, M.E., Bangs, M.J., Manguin, S., Coetzee, M., Mbogo, C.M., Hemingway, J., Patil, A.P., Temperley, W.H., Gething, P.W., Kabaria, C.W., Okara, R.M., Van Boekel, T., Godfray, H.C.J., Harbach, R.E. & Hay, S.I. (2010). The dominant Anopheles vectors of human malaria in Africa, Europe and the Middle East: occurrence data, distribution maps and bionomic précis. Parasites & Vectors, 3, 117. https://doi.org/10.1186/1756-3305-3-117

Skovgård, H. & Nachman, G. (2017). Modeling the temperature-and age-dependent survival, development, and oviposition rates of stable flies (*Stomoxys calcitrans*) (Diptera: Muscidae). Environmental Entomology, 46, 1130–1142. https://doi.org/10.1093/ee/nvx118

Tasioudi, K.E., Antoniou, S.E., Iliadou, P., Sachpatzidis, A., Plevraki, E., Agianniotaki, E.I., Fouki, C., Mangana-Vougiouka, O., Chondrokouki, E. & Dile, C. (2016). Emergence of lumpy skin disease in Greece, 2015. Transboundary and Emerging Diseases, 63, 260–265. https://doi.org/10.1111/tbed.12497

Taylor, D.B., Moon, R.D. & Mark, D.R. (2012). Economic impact of stable flies (Diptera: Muscidae) on dairy and beef cattle production. Journal of Medical Entomology, 49, 198–209. https://doi.org/10.1603/ME10050

Tuppurainen, E.S.M., Venter, E.H. & Coetzer, J.A.W. (2005). The detection of lumpy skin disease virus in samples of experimentally infected cattle using different diagnostic techniques. Onderstepoort Journal of Veterinary Research, 72, 153–164. https://doi.org/10.4102/ojvr.v72i2.213

Tuppurainen, E.S. & Oura, C.A.L. (2012). Lumpy skin disease: an emerging threat to Europe, the Middle East and Africa. Transboundary and Emerging Diseases, 59, 40–48. https://doi.org/10.1111/j.1865-1682.2011.01242.x

Tuppurainen, E.S., Lubinga, J.C., Stoltsz, W.H., Troskie, M., Carpenter, S.T., Venter, E.H. & Oura, C.A. (2013). Mechanical transmission of lumpy skin disease virus by *Rhipicephalus appendiculatus* male ticks. Epidemiology and Infection, 141, 425–430. https://doi.org/10.1017/S0950268812000805

Weiss, K.E. (1968). Lumpy skin disease. In: Virology Monographs 3 (pp. 111–131). Vienna: Springer-Verlag. https://doi.org/10.1007/978-3-662-39771-8_3

