## Supporting Information for "Using the basic reproduction number to assess the risk of transmission of lumpy skin disease virus by biting insects"

Simon Gubbins

*The Pirbright Institute, Ash Road, Pirbright, Surrey GU24 0NF, U.K.*

**Text S1	Deriving the basic reproduction number for lumpy skin disease virus**

***S1.1	Transmission model for LSDV***

To derive the basic reproduction number for lumpy skin disease virus (LSDV) in cattle, a mathematical model was developed to describe the transmission of LSDV between cattle via the bites of insect vectors.

The cattle population is assumed to be constant (*H*), except for disease-associated mortality, and is subdivided into the number of susceptible (i.e. uninfected), latent (i.e. infected, but not yet infectious), infectious and recovered animals, denoted by *S*, *E*, *I* and *R*, respectively. To allow for more general (gamma) distributions for the latent and infectious periods, the latent and infectious host populations, *E* and *I*, are subdivided into *n_i_* stages^[1]^ each of mean duration 1/*n_i_r_i_* (so that the latent and infectious periods follow gamma distributions with means 1/*r_i_* and variances 1/*n_i_r_i_*^2^ for *i*=*E*,*I*, respectively) (Anderson & Watson, 1980).

The vector population (*N*) is subdivided into the number of susceptible (i.e. uninfected) and infected insects, denoted by *X* and *Y*, respectively. Because transmission is mechanical and there is no need for the virus to replicate in the insect before transmission can occur, a vector becomes infectious as soon as it takes an infected blood meal (i.e. there is no extrinsic incubation period) and remains infectious until the virus is inactivated (at which point it become susceptible again) or the insect dies. Non-blood feeding individuals (e.g. adult male mosquitoes and biting midges or immature (egg, larval and pupal) life stages) are not considered in the model as they do not blood-feed and, hence, do not transmit LSDV.

The dynamics of infection in cattle are described by the following set of linked ordinary differential equations,

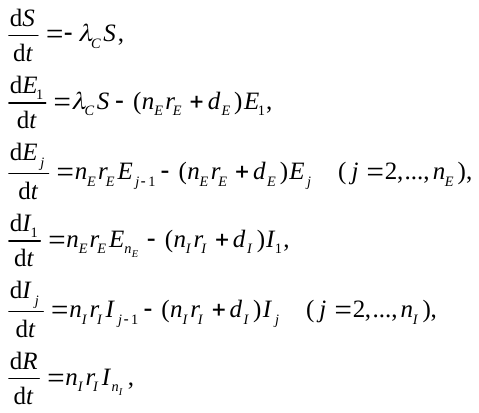

where *d_E_* and *d_I_* are the disease-associated mortality rates during the latent and infectious periods, respectively, and the force of infection for cattle is given by,

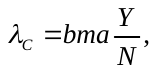

where *b* is the probability of transmission from insect to bovine, *m*=*N*/*H* is the vector-to-host ratio, *a* is the reciprocal of the time interval between blood meals and *Y*/*N* is the proportion of bites which are from infected insects.

The dynamics of infection in the insect vector are described by,

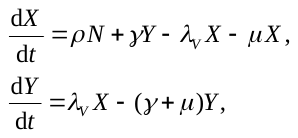

where *ρ* is the rate of recruitment to the biting population, *γ* is the virus inactivation rate and *μ* is the vector mortality rate. The force of infection for vectors is given by,

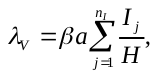

where *β* is the probability of transmission from bovine to insect and the remaining terms are defined above.

***S1.2	Basic reproduction number***

The basic reproduction number, *R*_0_, is calculated as the dominant eigenvalue, *r*(*K*), of the next-generation operator, *K* (Diekmann & Heesterbeek, 2000; Heffernan, Smith & Wahl, 2005). For the transmission model, -, the next generation operator is a matrix, **K**, whose elements (*K_ij_*) are the expected number of infected hosts or vectors (*i*) arising from a single infected host or vector (*j*). There is assumed to be no direct transmission between hosts or vectors (i.e. *K_HH_*=*K_VV_*=0), so that **K** can be written as,

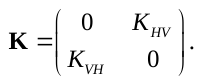

*Transmission from vector to host:* After an insect feeds on an infected bovine, it remains infected (and infectious) until the virus becomes inactivated or the insect dies, a period which lasts on average 1/(*γ*+*μ*) days, where *γ* is the virus inactivation rate and *μ* is the vector mortality rate. During this time the infected insect will bite susceptible cattle *a* times per day and a proportion, *b*, of these bites will result in a newly infected animal. Hence, the expected number of infected cattle per infected insect (*K_HV_*) is given by,

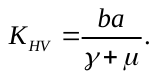

*Transmission from host to vector:* An infected bovine remains infectious for *D_I_* days, where

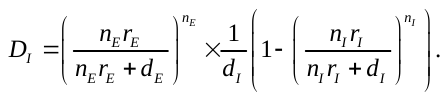

Here, the first term is the probability an infected bovine survives the latent period and the second term is the mean duration of infectiousness, allowing for disease associated mortality. During this time the infected bovine will be bitten by susceptible insects on average *m*×*a* times per day, a proportion, *β*, of which will result in a newly infected vector. Consequently, the expected number of infected insects per infected bovine (*K_VH_*) is given by,

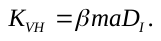

Some linear algebra shows that the dominant eigenvalue of **K** (i.e. *R*_0_) is,

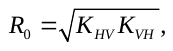

which, on substituting the expressions in and becomes,

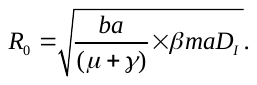

Mortality estimates for LSD outbreaks in the Balkans in 2015-2016 are very low, with the median within-herd mortality being zero (i.e. *d_E_*≈0 and *d_I_*≈0) in all affected countries (EFSA, 2017). In this case, the mean duration of infectiousness is the mean infectious period (i.e. *D_I_*=1/*r_I_*), which yields the expression for *R*_0_ in the main paper (equation (1)). The sensitivity of *R*_0_ estimates to this assumption is explored in Text S2.

**References for Text S1 (not included in the main paper)**

Anderson, D. & Watson, R. (1980). On the spread of a disease with gamma distributed latent and infectious periods. *Biometrika*, *67*, 191-198. <https://doi.org/10.1093/biomet/67.1.191>

Heffernan, J.M., Smith, R.J. & Wahl, L.M. (2005). Perspectives on the basic reproductive ratio. *Journal of the Royal Society Interface*, *2*, 281-293. <https://doi.org/10.1098/rsif.2005.0042>

**Text S2	Impact of disease-associated mortality on *R*_0_**

To assess its impact on the results, the uncertainty and sensitivity analyses were repeated including disease-associated mortality when computing the basic reproduction number (i.e. using equations and ). The level of mortality (*C*; i.e. the proportion of infected animals dying from disease) was sampled uniformly from the range 0 to 40%, which reflects the range reported in the field (Tuppurainen & Oura, 2012). The corresponding disease-associated mortality rates (*d_E_* and *d_I_*) were computed using the equation,

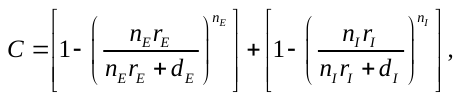

where the first term gives the proportion of infected animals dying during the latent period and the second the proportion dying during the infectious period. Two possibilities were considered: (i) mortality acts only during the infectious period (*d_E_*=0, *d_I_*>0); and (ii) mortality acts equally during the latent and infectious periods (*d_E_*=*d_I_*=*d*>0). Because calculation of the mortality rates requires them, the shape parameter (*n_E_*) and mean (1/*r_E_*) for the latent period and shape parameter (*n_I_*) for the infectious period were also included in the uncertainty and sensitivity analyses, and these were sampled from their marginal posterior distributions.

The uncertainty analyses indicated that the estimates for *R*_0_ for all five insect species are reduced when disease-associated mortality is included in the model (Table S3). However, the reduction in the median is small and that in the 95% prediction interval even smaller. The partial rank correlation coefficients (PRCCs) for the seven parameters included in the uncertainty and sensitivity analyses in the main paper (*b*, *β*, *a*, *m*, 1/*r*, *γ* and *μ*) did not change appreciably. In addition, the PRCCs for the additional parameters (i.e. *C*, *n_E_*, 1/*r_E_* and *n_I_*) were all close to zero, indicating they do not greatly influence the magnitude of *R*_0_. Accordingly, including disease-associated mortality over the range reported for LSDV in the field does not change the conclusions presented in the main paper.

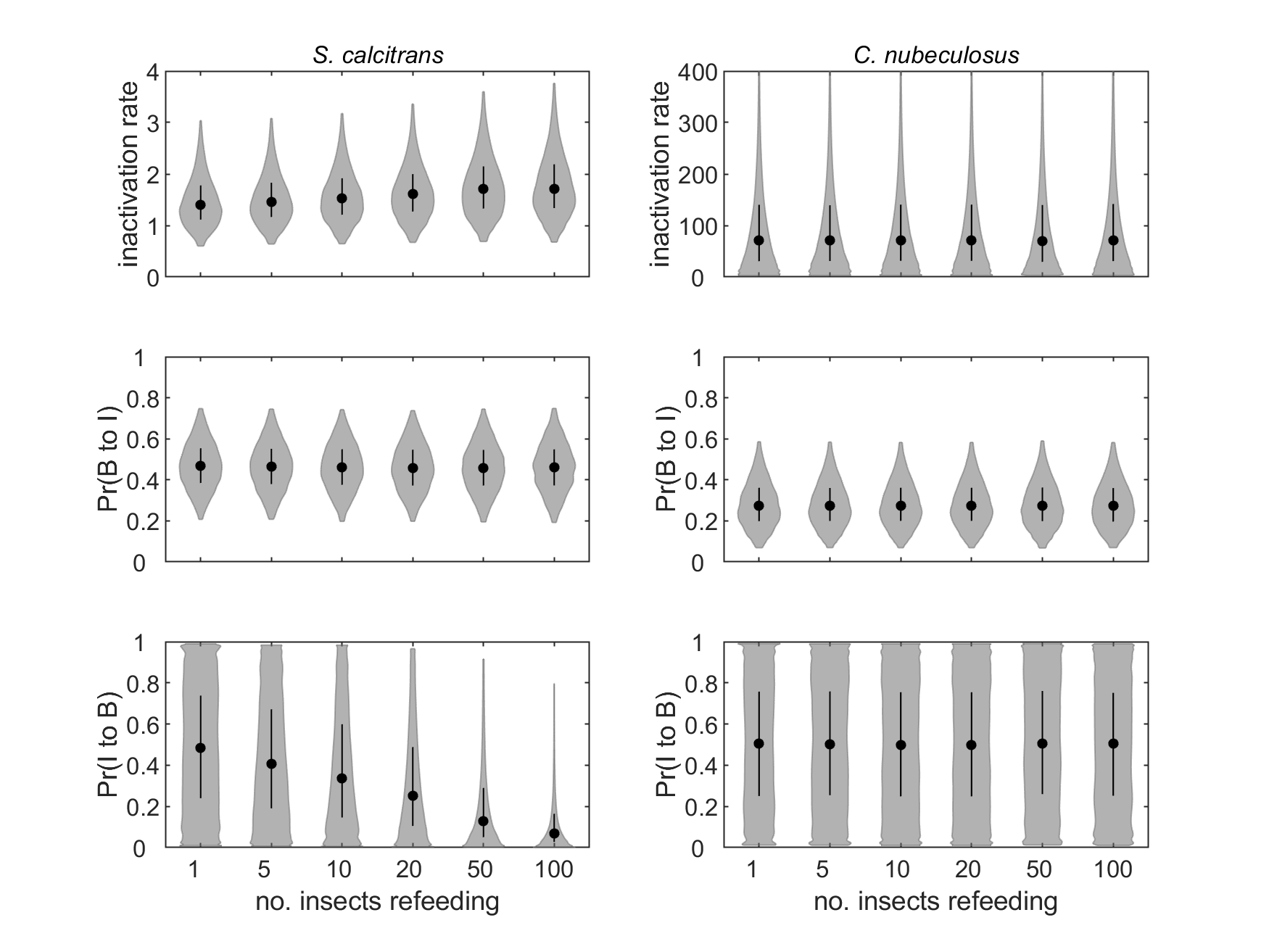

**Figure S1.** Effect of assumed number of insects refeeding on estimated parameters for mechanical transmission of lumpy skin disease virus by *Stomoxys calcitrans* (left column) and *Culicoides nubeculosus* (right column): virus inactivation rate (top row); probability of transmission from bovine to insect, Pr(B to I) (middle row); and probability of transmission from insect to bovine, Pr(I to B) (bottom row). Each plot shows the median (black circle), 25th and 75th percentiles (black line) and density (shape) for the marginal posterior distribution.

**Table S1.** Published data on the mechanical transmission of lumpy skin disease virus by biting insects (taken from Chihota et al., 2001, 2003).

(Key: *Y* - number of viral DNA positive insects; *N* - number of insects tested; *n* - number of insects that refed when attempting transmission to cattle; *I* - indicator of whether (*I*=1) or not (*I*=0) transmission occurred following refeeding)

| days post feeding | *S. calcitrans* | | | | *C. nubeculosus* | | | | *Ae. aegypti* | | | | *An. stephensi* | | | | *Cx. quinquefasciatus* | | | |
| --- | --- | --- | --- | --- | --- | --- | --- | --- | --- | --- | --- | --- | --- | --- | --- | --- | --- | --- | --- | --- |
|  | *Y* | *N* | *n*† | *I* | *Y* | *N* | *n*† | *I* | *Y* | *N* | *n*‡ | *I* | *Y* | *N* | *n*‡ | *I* | *Y* | *N* | *n*‡ | *I* |
| 0 | 5 | 12 | - | - | 3 | 12 | - | - | 5 | 5 | - | - | 6 | 10 | - | - | 7 | 10 | - | - |
| 1 | 3 | 12 | - | 0 | 0 | 12 | - | - | - | - | - | - | - | - | - | - | - | - | - | - |
| 2 | 0 | 12 | - | 0 | 0 | 12 | - | - | 3 | 5 | 200 | 1§ | 4 | 10 | - | - | 5 | 10 | - | - |
| 3 | 0 | 12 | - | 0 | 0 | 12 | - | 0 | 4 | 5 | 200 | 1 | - | - | - | - | - | - | - | - |
| 4 | 0 | 12 | - | - | 0 | 12 | - | 0 | 4 | 5 | 200 | 1 | 3 | 10 | 60 | 0§ | 3 | 10 | 10 | 0¶ |
| 5 | 0 | 12 | - | - | 0 | 12 | - | 0 | 2 | 5 | 200 | 1 | - | - | - | - | - | - | - | - |
| 6 | 0 | 12 | - | - | 0 | 12 | - | - | 3 | 5 | 200 | 1 | 2 | 10 | - | - | 1 | 10 | - | - |
| 7 | 0 | 12 | - | - | 0 | 12 | - | - | - | - | - | - | - | - | - | - | - | - | 10 | 0§ |
| 8 | 0 | 12 | - | - | 0 | 12 | - | - | - | - | - | - | 1 | 10 | - | - | 0 | 5 | - | - |
| 9 | 0 | 10 | - | - | 0 | 12 | - | - | - | - | - | - | - | - | - | - | - | - | - | - |
| 10 | 0 | 8 | - | - | 0 | 7 | - | - | - | - | - | - | 0 | 6 | - | - | 0 | 6 | - | - |
| 20 | 0 | 8 | - | - | 0 | 3 | - | - | - | - | - | - | 0 | 3 | - | - | 0 | 4 | - | - |

† number of insects refeeding not specified; parameters estimated assuming *n*=1, 5, 10, 20, 50 or 100.

‡ approximate number of insects refed

§ transmission was attempted to two calves (and the outcome was the same for both animals)

¶ transmission was attempted to four calves (and the outcome was the same for all animals)

**Table S2.** Detection of lumpy skin disease virus in experimentally infected cattle.

(Key: *t_LN_* - time (post inoculation) of last negative sample; *t_FP_* - time of first positive sample; *t_LP_* - time of last positive sample; *t_FN_* - time of first subsequent negative sample)

| study | ID | blood (PCR) | | | | blood (VI) | | | | skin lesions | | | |
| --- | --- | --- | --- | --- | --- | --- | --- | --- | --- | --- | --- | --- | --- |
|  |  | *t_LN_* | *t_FP_* | *t_LP_* | *t_FN_* | *t_LN_* | *t_FP_* | *t_LP_* | *t_FN_* | *t_LN_* | *t_FP_* | *t_LP_* | *t_FN_* |
| Chihota et al. 2001 | TU39 | 5 | 6 | - | - | - | - | - | - | - | - | - | - |
|  | TW6 | 7 | 8 | - | - | - | - | - | - | - | - | - | - |
|  | TW7 | 6 | 7 | - | - | - | - | - | - | - | - | - | - |
|  | TW8 | 8 | 9 | - | - | - | - | - | - | - | - | - | - |
|  | TW9 | 9 | 10 | - | - | - | - | - | - | - | - | - | - |
|  | TW10 | 9 | 10 | - | - | - | - | - | - | - | - | - | - |
| Tuppurainen et al. 2005 | 1 | 1 | 2 | 24 | 25 | 5 | 6 | 16 | 17 | 5 | 6† | 33‡ | 39‡ |
|  | 2 | 1 | 2 | 28 | 29 | 5 | 6 | 16 | 17 | 6 | 7† | 33‡ | 39‡ |
|  | 3 | 3 | 4 | 27 | 28 | 10 | 11 | 16 | 17 | - | - | - | - |
|  | 4 | 2 | 3 | 26 | 27 | 4 | 5 | 16 | 17 | - | - | - | - |
|  | 7 | 2 | 3 | 13 | 14 | 15 | 16 | 16 | 17 | 9 | 10† | 25§ | 26§ |
|  | 8 | 8 | 9 | 26 | 27 | 4 | 5 | 16 | 17 | 7 | 8† | 18§ | 25§ |
| Babiuk et al. 2008 | 4 | 3 | 6 | 9 | 12 | - | - | - | - | - | - | - | - |
|  | 5 | 6 | 9 | 15 | - | - | - | - | - | - | - | - | - |
|  | 7 | 6 | 9 | 9 | 12 | - | - | - | - | - | - | - | - |
|  | 8 | 3 | 6 | 15 | 18 | 6 | 9 | 9 | 12 | - | - | - | - |

† time of first appearance of skin lesions (biopsy samples not taken until 10 days post inoculation)

‡ based on detection of virus in biopsy samples by transmission electron microscopy

§ based on detection of viral DNA in biopsy samples by PCR

**Table S3.** Basic reproduction number† (*R*_0_) for lumpy skin disease virus (LSDV) for five species of biting insect under different assumptions about disease-associated mortality.

| species | proxy measure of infectiousness | | |
| --- | --- | --- | --- |
|  | skin biopsies | blood (PCR) | blood (virus isolation) |
| no disease-associated mortality | |  |  |
| *S. calcitrans*‡ | 15.5 (1.4, 81.9) | 13.1 (1.2, 70.8) | 9.7 (0.9, 51.7) |
| *C. nubeculosus*‡ | 1.8 (0.06, 13.5) | 1.5 (0.05, 11.3) | 1.1 (0.04, 8.8) |
| *Ae. aegypti* | 7.4 (1.3, 17.6) | 6.3 (1.1, 14.5) | 4.6 (0.8, 11.2) |
| *An. stephensi* | 1.6 (0.2, 6.0) | 1.4 (0.2, 5.0) | 1.0 (0.1, 3.9) |
| *Cx. quinquefasciatus* | 0.8 (0.09, 3.5) | 0.7 (0.08, 2.8) | 0.5 (0.06, 2.2) |
| mortality during infectious period only | |  |  |
| *S. calcitrans*‡ | 14.8 (1.3, 75.3) | 12.0 (1.1, 62.4) | 9.0 (0.8, 47.9) |
| *C. nubeculosus*‡ | 1.7 (0.05, 12.4) | 1.4 (0.05, 10.4) | 1.0 (0.03, 7.9) |
| *Ae. aegypti* | 7.0 (1.2, 16.4) | 5.8 (1.0, 13.8) | 4.3 (0.7, 10.4) |
| *An. stephensi* | 1.5 (0.2, 5.7) | 1.3 (0.2, 4.8) | 0.9 (0.1, 3.6) |
| *Cx. quinquefasciatus* | 0.8 (0.09, 3.2) | 0.6 (0.07, 2.7) | 0.5 (0.05, 2.0) |
| equal mortality during latent and infectious periods | | |  |
| *S. calcitrans*‡ | 14.4 (1.3, 75.6) | 12.0 (1.1, 64.9) | 8.7 (0.8, 46.0) |
| *C. nubeculosus*‡ | 1.7 (0.05, 12.8) | 1.4 (0.04, 10.1) | 1.0 (0.03, 7.9) |
| *Ae. aegypti* | 6.9 (1.2, 16.5) | 5.7 (1.0, 13.7) | 4.2 (0.7, 10.3) |
| *An. stephensi* | 1.5 (0.2, 5.7) | 1.2 (0.2, 4.6) | 0.9 (0.1, 3.6) |
| *Cx. quinquefasciatus* | 0.8 (0.09, 3.2) | 0.6 (0.07, 2.6) | 0.5 (0.05, 2.0) |

† median (95% prediction interval) based on replicated Latin hypercube sampling

‡ estimates are shown for the analysis assuming 100 insects refed when attempting transmission to cattle

1. The number of stages is equivalent to the shape parameter for a gamma distribution, but is constrained to take integer values.
